# Taking the sub-lexical route: brain dynamics of reading in the semantic variant of Primary Progressive Aphasia

**DOI:** 10.1101/847798

**Authors:** V. Borghesani, L.B.N. Hinkley, K. G. Ranasinghe, M. M. C. Thompson, W. Shwe, D. Mizuiri, M. Lauricella, E. Europa, S. Honma, Z. Miller, B. Miller, K. Vossel, M. Henry, J. F. Houde, M.L. Gorno-Tempini, S. S. Nagarajan

## Abstract

Reading aloud requires mapping an orthographic form to a phonological one. The mapping process relies on sub-lexical statistical regularities (e.g., “oo” to |u□|) or on learned lexical associations between a specific visual form and a series of sounds (e.g., *yacht* to /j□t/). Computational, neuroimaging, and neuropsychological evidence suggest that sub-lexical, phonological and lexico-semantic processes rely on partially distinct neural substrates: a dorsal (occipito-parietal) and a ventral (occipito-temporal) route, respectively.

Here, we investigated the spatiotemporal features of orthography-to-phonology mapping, capitalizing on the time resolution of magnetoencephalography and the unique clinical model offered by patients with semantic variant of Primary Progressive Aphasia (svPPA). Behaviorally, svPPA patients manifest marked lexico-semantic impairments including difficulties in reading words with exceptional orthographic to phonological correspondence (irregular words). Moreover, they present with focal neurodegeneration in the anterior temporal lobe (ATL), affecting primarily the ventral, occipito-temporal, lexical route. Therefore, this clinical population allows for testing of specific hypotheses on the neural implementation of the dual-route model for reading, such as whether damage to one route can be compensated by over-reliance on the other. To this end, we reconstructed and analyzed time-resolved whole-brain activity in 12 svPPA patients and 12 healthy age-matched controls while reading irregular words (e.g., *yacht*) and pseudowords (e.g., *pook*).

Consistent with previous findings that the dorsal route is involved in sub-lexical, phonological processes, in control participants we observed enhanced neural activity over dorsal occipito-parietal cortices for pseudowords, when compared to irregular words. This activation was manifested in the beta-band (12-30 Hz), ramping up slowly over 500 ms after stimulus onset and peaking at ∼800 ms, around response selection and production. Consistent with our prediction, svPPA patients did not exhibit this temporal pattern of neural activity observed in controls this contrast. Furthermore, a direct comparison of neural activity between patients and controls revealed a dorsal spatiotemporal cluster during irregular word reading. These findings suggest that the sub-lexical/phonological route is involved in processing both irregular and pseudowords in svPPA.

Together these results provide further evidence supporting a dual-route model for reading aloud mediated by the interplay between lexico-semantic and sub-lexical/phonological neuro-cognitive systems. When the ventral route is damaged, as in the case of neurodegeneration affecting the ATL, partial compensation appears to be possible by over-recruitment of the slower, serial attention-dependent, dorsal one.

**Abbreviated Summary:** Borghesani et al. investigate brain dynamics during irregular word reading using magnetoencephalographic imaging in patients with semantic variant of primary progressive aphasia. Due to ventral anterior temporal lobe neurodegeneration, patients show greater reliance of dorsal, occipito-parietal brain regions – providing novel evidence for the interplay between ventral and dorsal routes for reading.

## Introduction

Reading aloud, the process by which a visual input is translated into an auditory output, entails a series of steps involving different neural systems: from vision to motor control. Crucially, this requires mapping an orthographic form to its corresponding phonological form. Language-specific statistical regularities allow readers to spell out words never encountered before (sometimes referred to as pseudowords or nonwords, e.g., *pook*) by selecting the most plausible phonological representations. In non-transparent languages such as English, access to meaning is necessary to correctly pronounce words with exceptional spelling-to-sound correspondence (i.e., irregular words, e.g., *yacht*). Computational models of reading assume that this process involves both storage of the relationship between orthography and phonology (sub-lexical, phonological representations), as well as the knowledge of learned words (lexico-semantic representations) (Perry *et al.*, 2007). For instance, the dual-route cascaded model (DRC), explicitly assumes the interplay of two computationally distinct routes: a sub-lexical and a lexical one, needed to select the appropriate phonological output for pseudowords and irregular words respectively (Coltheart *et al.*, 1993, 2001). Alternatively, a single system with interactive orthographic, phonological, and semantic representations has been posited and tested computationally - the so-called “triangle” model (Plaut *et al.,* 1996). The original formulations of these accounts diverged on the nature of the orthography-to-phonology processes (i.e., are sub-word representations required or could the mapping happen at the whole word level?), and on whether they postulated the independence of lexical from semantic processes (Taylor *et al.*, 2013). In their latest instantiations, all predominant cognitive models agree that reading requires the interplay of different representations (i.e., sub-lexical/phonological vs. lexico-semantic), with different stimuli (and tasks) shifting the emphasis towards one or the other.

The idea that phonological and lexico-semantic representations are linked to (at least) two different cognitive mechanisms supported by (at least) two distinct neural substrates originates in the neuropsychological evidence of a double dissociation, as two kinds of acquired dyslexia have been identified (Marshall and Newcombe, 1973) - phonological and surface dyslexia. Phonological dyslexia is characterized by a deficit in reading pseudowords, with relatively preserved reading of irregular words, suggesting a selective disruption of the grapheme-to-phoneme conversion route (Coltheart, 2006). Typical errors in phonological dyslexia are called lexicalizations, where pseudowords are read as real words: e.g., *pook* read as *hook*. In contrast, surface dyslexia is characterized by relatively spared reading of pseudowords with impaired reading of irregular words. Typical errors in surface dyslexia are called regularizations: e.g., *yacht* read as /jæt□t/, instead of /j□t/ (Coltheart, 2006), where irregular words are read as phonologically-plausible pseudowords. This profile suggests an impairment of the lexico-semantic system (Woollams *et al.*, 2007).

Neuroimaging studies provide complementary observations indicating that sub-lexical and lexical processes rely on partially distinct neural substrates. Sub-lexical processes have been associated with dorsal structures in the inferior parietal lobe (i.e., supramarginal gyrus, angular gyrus, and temporoparietal junction) and the posterior inferior frontal gyrus (IFG) by studies investigating pseudoword reading, i.e., regular orthographic-to-phonological mappings (Jobard *et al.*, 2003; Mechelli *et al.*, 2003; Graves *et al.*, 2010; Taylor *et al.*, 2013; Sliwinska *et al.*, 2015). In contrast, lexical processes, have proven harder to isolate, but have been linked to ventral brain structures in the temporal lobe, especially its anterior portion (ATL), which appears to be involved in semantic-mediated reading of low-frequency irregular words (Graves *et al.*, 2010; Wilson *et al.*, 2012; Taylor *et al.*, 2013; Hoffman *et al.*, 2015; Provost *et al.*, 2016).

Computational models and behavioral evidence suggest key computational differences between the two routes: while activated simultaneously for all word-like inputs, only the lexical one can lead to the correct pronunciation of irregular words, while only the sub-lexical one allows reading of pseudowords. To date however, little is known about the neural dynamics characterizing these processes, although different temporal profiles have been suggested. On one hand, serial effects have been associated with the sub-lexical route, arguably indicating a slow grapheme-by-grapheme conversion mechanism (e.g., Weekes, 1997). On the other hand, it has been suggested that the ventral route operates in parallel, within a fast, automatic spreading of activation (Coltheart *et al.*, 2001). Hence, divergence of the two routes should be detected in a relatively late time frame, after ∼400 ms, close to response selection and production. Many reading studies have focused on event-related potential (ERPs), which capture stimulus-locked, phased-locked evoked activity immediately following the presentation of written stimuli (for a review, Salmelin, 2007; Grainger and Holcomb, 2009). These studies may have overlooked slower, later, not necessarily phased-locked effects, which can be better captured by analyses of task-induced neural oscillatory changes.

While neuroimaging studies can provide only correlational evidence, neuromodulation techniques and neuropsychological findings have strengthened the evidence of a neural dissociation between lexical and sub-lexical computations. For instance, recent evidence that repetitive transcranial magnetic stimulation (rTMS) over left ventral ATL leads to regularization errors has corroborated the critical role played by this region in irregular words reading (Ueno *et al.*, 2018). In this context, patients with semantic variant of Primary Progressive Aphasia (svPPA) are an ideal clinical model for testing hypotheses on the interplay between dorsal (parietal) vs. ventral (temporal) routes. SvPPA is associated with (relatively) focal atrophy of the ATL which, in time, spreads towards the orbitofrontal cortex (Galton *et al.*, 2001; Rosen *et al.*, 2002, Brambati *et al.*, 2009*b*). Clinically, svPPA patients present with fluent speech and pervasive semantic memory deficits (Hodges *et al.*, 1992). It is well established that patients with svPPA manifest surface dyslexia (Hodges *et al.*, 1992, 1999; Neary *et al.*, 1998) with a clear effect of word frequency and length on performance accuracy in irregular word reading (Patterson and Hodges, 1992; Jefferies *et al.*, 2004; Patterson *et al.*, 2006). In particular, svPPA patients produce so-called over-regularization errors, such as reading “*sew”* as “*sue”*, i.e., the application of legitimate alternative correspondence between graphemes and phonemes (Woollams *et al.*, 2007; Wilson *et al.*, 2009). Crucially, the severity of the reading deficit has been linked to the extent of the semantic loss (Patterson and Hodges, 1992; Graham *et al.*, 2000; Jefferies *et al.*, 2004). Finally, irregular word reading accuracy has been directly linked to the integrity of left ATL (vs. pseudowords to left temporo-parietal structures, Brambati *et al.*, 2009*a*). To date, only one study has attempt to investigate the correlates of surface dyslexia in svPPA with a neuroimaging experiment: it has been suggested that, given the damage to ATL, svPPA patients recruit the intraparietal sulcus to read irregular words as if they were pseudowords (Wilson *et al.*, 2009).

Here, we investigate the spatiotemporal dynamics of reading capitalizing on the high temporal resolution offered by magnetoencephalographic imaging, and on the known clinical and neuro-anatomical features of svPPA. Our approach, compatible with both prominent cognitive theories on reading, enables us to directly tackle the neural correlates of the interplay between ventral and dorsal routes. We hypothesized that svPPA patients, given their ATL atrophy and subsequent lexical-semantic deficit, would rely more on sub-lexical/phonological processes to read irregular words, subserved by the relatively structurally intact dorsal, occipito-parietal cortices. Specifically, we predict an over-recruitment of the dorsal route during reading of irregular words in svPPA patients. Given the posited serial and slower nature of the dorsal route computation, we expect this over-recruitment to occur at later latencies.

## Materials and methods

### Subjects

Twelve svPPA patients (8 female, 67.92 ± 7.33 years old) and 12 healthy controls matched for age, education, and gender (9 female, 72.17 ± 4.04 years old) were recruited through the University of California San Francisco (UCSF) Memory and Aging Center (MAC). All participants were right-handed native English speakers, had no history of developmental dyslexia, and no contraindications to MEG. Patients met currently published diagnostic criteria for svPPA, as determined by a team of clinicians and physicians, based on a detailed medical history, comprehensive neurological assessment, and standardized neuropsychological and language evaluations (Kramer *et al.*, 2003; Gorno-tempini *et al.*, 2011). All patients included in the study scored at least 15 out of 30 on the Mini-Mental Status Exam (MMSE) and were sufficiently functional to be scanned. In addition, each patient had left-predominant (rather than right) anterior temporal lobe atrophy to maximize our chances of detecting surface dyslexia in a homogenous sample (Binney *et al.*, 2016). Demographic information and neuropsychological data are shown in Table 1. Two-sample t-tests (two-tailed distributions, significance threshold set at p<0.05) were used to statistically assess group differences between these svPPA patients and healthy controls (using published data; Gorno-Tempini *et al.*, 2004; Binney *et al.*, 2016; Watson *et al.*, 2018). Approved study protocols by the UCSF Committee on Human Research were followed, and all subjects provided written informed consent.

**Table 1.**
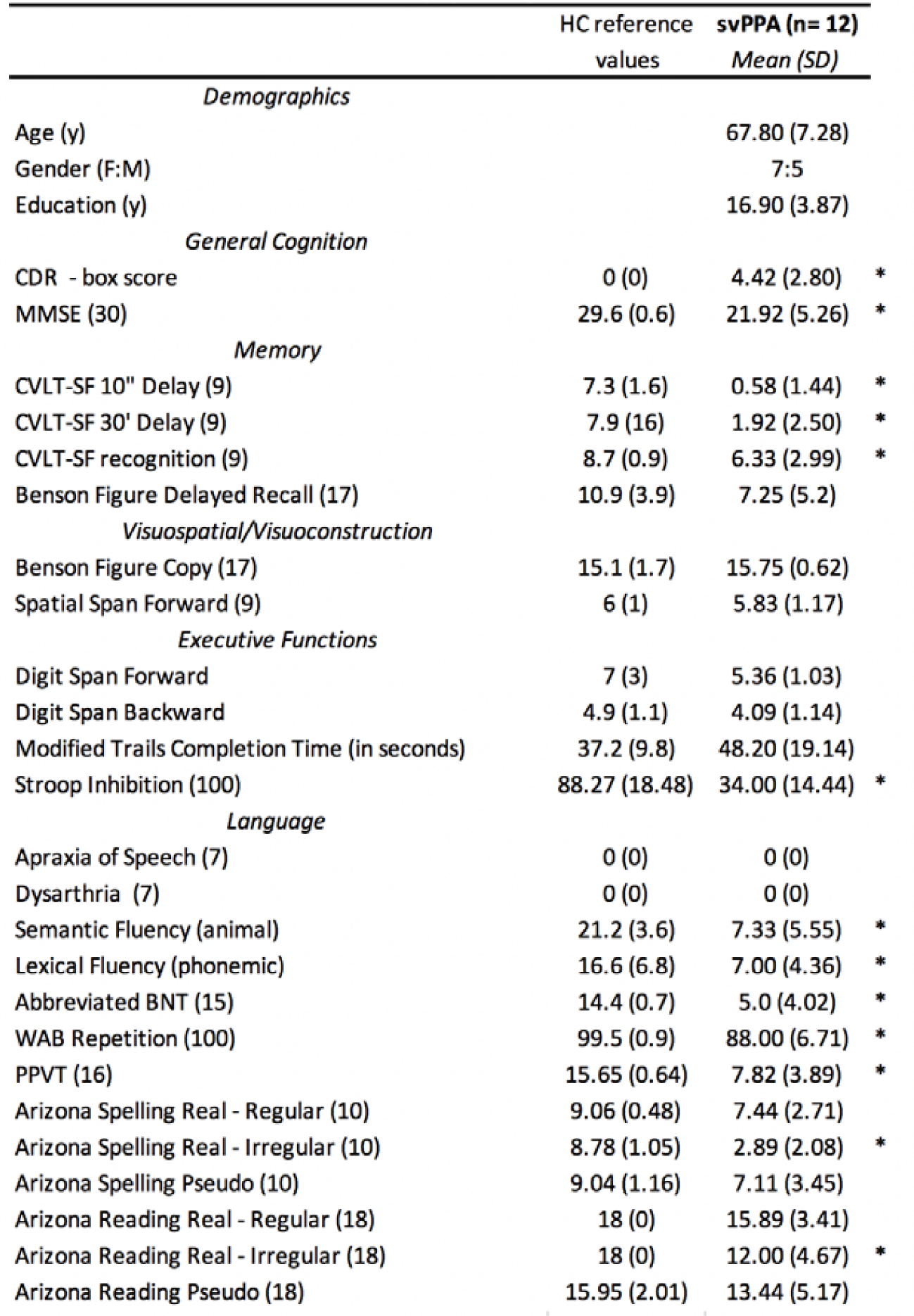
Demographic and neuropsychological profile of svPPA patients. All participants were right handed native English speakers. The patient group includes 12 cases of semantic variant of Primary Progressive Aphasia (svPPA). Scores shown are mean (standard deviation). *indicate values significantly different from controls (p<0.05). CDR = Clinical Dementia Rating, MMSE = Mini-Mental State Exam. CVLT=California Verbal Learning Test, BNT = Boston Naming Test, WAB = Western Aphasia Battery, PPVT = Picture Vocabulary Test.

### Stimuli and Experimental Design

Visual stimuli were projected into the magnetically shielded MEG scanner room. Participants were instructed to read aloud the string of letters immediately following its appearance on the screen, speaking their responses into a MEG-compatible microphone. The stimuli consisted of 40 regular words, i.e., words with regular grapheme-to-phoneme correspondence (e.g., *fact*), 40 irregular words, i.e., words with inconsistent grapheme-to-phoneme correspondence (e.g., *choir*), and 70 pronounceable strings of letters with no semantic representation (e.g., *pook*). Real words and pseudowords (n=20) were drawn from the Arizona reading list (Beeson and Rising, 2010). An additional 50 pseudowords were generated using an established computerized multi-lingual pseudo-word generator Wuggy (Keuleers and Brysbaert, 2010). Three sets of 50 pseudowords each were generated by submitting a list of 50 real word nouns to the multilingual pseudoword generator using the English language setting for sub-syllabic structures and transition frequencies. The first 50 pseudowords set was checked by a native speaker informant, who suggested no replacement from the backup second or third sets. Examples of stimuli from each category, along with key psycholinguistic variables are shown in Table 2. Words’ number of letters, frequency (i.e., log-transformation of the Thorndike-Lorge written frequency), age of acquisition, concreteness, familiarity, and imaginability were extracted from the MRC Psycholinguistic Database (http://websites.psychology.uwa.edu.au/school/MRCDatabase/uwa_mrc.htm). Bigram frequency, (total bigram count), number of orthographic and phonological neighbors, as well as orthography-phonology consistency were derived from The English lexicon project database (Balota et al., 2007; https://elexicon.wustl.edu). Orthography-Phonology Consistency (OPC), quantifying the phonological relatedness between a word and its orthographic relatives, was calculate by dividing the number of orthographic neighbors that share the same pronunciation (e.g., for *north, forth*) by the total number of orthographic neighbors (e.g., for *north, worth* and *forth*). Orthography-Semantics Consistency (OSC), quantifying the semantic relatedness between a word and its orthographic relatives, was derived from the Orthography-Semantics Consistency Database (Marelli and Amenta, 2018; http://www.marcomarelli.net/resources). None of these variables was statistically different between regular and irregular words, except for orthography-phonology consistency: regular words have a significantly higher ratio of orthographic neighbors that share the same pronunciation (p<0.0001). Pseudowords did not statistically differ from the full set of words in terms of number of letters, number of orthographic neighbors, and bigram frequency. However, the difference in the number of orthographic neighbors between pseudowords and irregular words reached significance (p=0.044). Finally, it should be noted that using other frequency measures such as the KF frequency (Kucera & Francis, 1967), the HAL frequency (Hyperspace Analogue to Language; Lund & Burgess, 1996), or the movie subtitles frequency as reported in The English lexicon project database leads to the same results.

**Table 2.** Psycholinguistic characteristics of the stimuli. All words (n=80) and a portion of the pseudowords (n=20) were taken from the Arizona reading list. An additional set of 50 pseudowords were generated with Wuggy. Values shown are mean (standard deviation), * denotes a significant difference between regular and irregular words, while # denotes a significant difference between pseudowords and irregular words. Words frequency, age of acquisition, concreteness, familiarity, and imaginability values were extracted from the MRC Psycholinguistic Database. Number of orthographic neighbors, bigram frequency (total bigram count), and orthography-phonology consistency (OPC) were derived from the English Lexicon Project Database. The orthography-semantics consistency (OSC) was derived from the Orthography-Semantics Consistency Database.

| **TABLE 2. Stimuli Examples and Psycholinguistic Variables** |  |  |  |
| --- | --- | --- | --- |
|  | **Regular Words** | **Irregular Words** | **Pseudo Words** |
|  | n = 40 | n = 40 | n = 20 + 50 |
|  | *Mean (SD)* | *Mean (SD)* | *Mean (SD)* |
| Examples | *fact, bribe, magnet* | *choir, castle, yacht* | *pook, andon, codle* |
| Length (# of letters) | 5.1 (0.91, 4 to 7) | 5.1 (0.91, 4 to 7) | 4.8 (0.99, 3 to 8) |
| Orthographic Neighbors | 6.15 (5.99) | 4.48 (4.57) | 6.64 (5.69) # |
| Bigram Frequency | 15657.10 (7009.51) | 13349.93 (8778.66) | 14016.77 (8324.17) |
| Phonological Neighbors | 13.15 (10.40) | 15.35 (12.79) | n/a |
| Frequency | 2.43 (0.61) | 2.39 (0.70) | n/a |
| Age of Acquisition | 308.43 (28.57) | 315.75 (78.97) | n/a |
| Concreteness | 469.15 (107.07) | 476.21 (105.74) | n/a |
| Familiarity | 551.93 (31.52) | 530.27 (56.33) | n/a |
| Imaginability | 500.46 (95.35) | 485.53 (106.30) | n/a |
| OPC | 0.65 (0.32) | 0.24 (0.34) * | n/a |
| OSC | 11.29 (1.85) | 11.65 (1.49) | n/a |

Words and pseudowords were presented in two separate runs, with words always preceding pseudowords. Words were presented twice each in a fixed, sequential order (1 second display, 1.7-2.1 second inter-stimulus interval (ISI)). Pseudowords were presented twice each in randomized order (2 second display, 1.7-2.1 second ISI). Vocal responses were digitized on separate analog-to-digital channels (ADCs), marked through amplitude threshold detection, and verified by hand through visual inspection manually in each dataset. Participants were instructed to read the stimulus as quickly and accurately as possible; however, if the patient needed more time **to** respond, the duration of the ISI was extended to ensure that responses were captured without interfering with the next trial.

### Behavioral data

Responses were recorded with Audacity (www.audacityteam.org) and coded offline by three independent raters as either correct, error, or no response. Errors with irregular words were further marked as regularizations if clearly influenced by the orthographic form of the word (e.g. reading of *shove* as to rhyme with *drove*). Similarly, errors with pseudowords were marked as lexicalizations if leading to the production of a real word (e.g. reading of *glope* as *globe*). Reaction times and accuracy (percentage of errors) were statistically analyzed using an analysis of variance (ANOVA) based on the three word types (regular, irregular, and pseudo-words) and cohorts (controls vs. svPPA patients) using the Python statistical library Statsmodels (www.statsmodels.org). Post-hoc *t-*tests were used to compare RTs and percentage accuracy of the two cohorts across the three word types. Similarly, differences in error types were analyzed with an ANOVA 2 (percentage of regularizations vs. percentage of lexicalizations) * 2 (controls vs. svPPA patients), and post-hoc t-tests performed to directly compare cohorts across error types. Explorative, post-hoc analyses of the effect of frequency and orthography-to-phonology consistency are reported in Suppl.Fig.1. Data from one outlier in the svPPA cohort were excluded from the behavioral analyses (across all conditions, the average percentage of error was 84.29 vs. 14.34 of the rest of the cohort, while the average reaction times as 1284.1 ms vs 888.8 ms in the rest of cohort).

### MRI protocol and analyses

Structural T1-weighted images were acquired on a 3T Siemens system (Siemens, Erlagen, Germany) at the UCSF Neuroscience Imaging Center, equipped with a standard quadrature head coil with sequences, previously described in (Mandelli et al., 2014). MRI scans were acquired within 1 year of the MEG data acquisition. To identify regions of atrophy, svPPA patients were compared to a separate set of 25 healthy controls collected using the same protocol (14 females, mean age 66.2 ± 8.5) using voxel-based morphometry (VBM). Image processing and statistical analyses were performed using the VBM8 Toolbox implemented in Statistical Parametric Mapping (SPM8, Wellcome Trust Center for Neuroimaging, London, UK, http://www.fil.ion.ucl.ac.uk/spm) running under Matlab R2013a (MathWorks). The images were segmented into grey matter, white matter, and CSF, bias corrected, and then registered to the Montreal Neurological Institute (MNI). The grey matter value in each voxel was multiplied by the Jacobian determinant derived from spatial normalization in order to preserve the total amount of grey matter from the original images. Finally, to ensure the data were normally distributed and to compensate for inexact spatial normalization, the modulated grey matter images were smoothed with a full-width at half-maximum (FWHM) Gaussian kernel filter of 8×8×8 mm. A general linear model (GLM) was then fit at each voxel, with one variable of interest (group), and three confounds of no interest: gender, age, education, and total intracranial volume (calculated by summing across the grey matter, white matter and CSF images). The resulting statistical parametric map (SPM) was thresholded at p<0.001 with family wise error (FWE) correction voxel-wise and used to visualize svPPA patients atrophy pattern (see Fig.1C), as well as build an exclusive mask for the MEG source reconstruction analyses (see below).

### MEG protocol and analyses

Neuromagnetic recordings were conducted using a whole-head 275 axial gradiometer MEG system (CTF, Coquitlam, BC, Canada) at a sampling rate of 1200 Hz. Head position was recorded before and after each block using three fiducial coils (nasion, left/right preauricular) placed on the participant. After data acquisition, each dataset was visually inspected to identify and remove noisy MEG sensors as well as trials with artifacts or missing responses that exceeded 1 pT fluctuations. Datasets were then epoched with respect to stimuli presentation onset (stimulus-locked trials) or vocal response onset (response-locked trials). In order to further remove excessive noisy artifacts (e.g., dental work, gritting) from some trials, datasets were cleaned using dual signal subspace projection (DSSP) and filtered between 4 and 115 Hz prior to source reconstruction (Sekihara *et al.*, 2016; Cai *et al.*, 2018).

We then reconstructed whole-brain oscillatory activity using the Neurodynamic Utility Toolbox for MEG (NUTMEG; http://nutmeg.berkeley.edu), which implements a time–frequency optimized adaptive spatial filtering technique to estimate spatiotemporal locations of neural sources. A tomographic volume of source locations was computed through an adaptive spatial filter (8 mm lead field) that weights each location relative to the signal of the MEG sensors (Dalal *et al.*, 2008). Data was reconstructed in the beta (12-30 Hz) range using partially overlapping time windows (200 ms, 50 ms step size) optimized for capturing induced, non-phase locked changes in the MEG signal generated by neural activity. Source power for each location was derived through a noise-corrected pseudo-F statistic expressed in logarithmic units (decibels) comparing signal magnitude during an active experimental time window (i.e., from 0, stimuli onset, to 900 ms) versus a baseline control window (i.e., 250 ms preceding stimuli onset) (Robinson and Vrba, 1999).

We focused our analyses on beta band (12-30 Hz) activations during reading of irregular words and pseudowords for three main reasons. First, we wished to overcome limitations of previous studies by looking not only at evoked but also induced changes in brain activity (i.e., modulations of ongoing oscillatory processes that are not phased-locked). This is especially relevant when the effects might occur later with respect to stimulus onset, as in the current setting, where differences between controls and patients in early, low-level visual processes are not expected, while divergence in later stages is predicted by both behavioral evidence and neuroanatomical findings. Furthermore, increased latency in response to linguistic stimuli has been observed in svPPA patients, and analyses of oscillatory behavior might be more sensitive than ERPs in detecting abnormalities in slow neural responses with increased trial-by-trial variability and thus decreased phase-locking (Kielar *et al.*, 2018). Second, we aimed at providing functional interpretation of our findings. The beta band has been extensively related to cortical activations. Specifically, desynchronization (i.e., beta band suppression) is a reliable indicator of heightened neural activity (Yuan *et al.*, 2010; Stevenson *et al.*, 2011) and covaries with BOLD signal in fMRI. Also, the beta band has been shown to capture predominantly left lateralized language functions, such as reading, and svPPA is a strongly left lateralized neurodegenerative disease (Wang *et al.*, 2012; Hinkley *et al.*, 2016). Moreover, beta band oscillations have been reliably associated with both memory and motor components of spoken word production (Herman *et al.*, 2013; Piai *et al.*, 2015). Third, in order to highlight the difference between the lexical and sub-lexical route, the key conditions that we compared were irregular words (as they can be read correctly only relying on lexical knowledge) and pseudowords (as they can be read only via sub-lexical processes). Regular words can be read using either route and thus do not offer any clear insight on the neural processes underlying lexical and sub-lexical processes.

Single-subject beamformer reconstructions were spatially normalized by applying each subject’s T1-weighted transformation matrix to their statistical map, and group analyses were performed with statistical nonparametric mapping (SnPM). From each voxel, we obtained a permuted distribution, then we estimated the significance of each pseudo-F value from its position in this permuted distribution (Singh *et al.*, 2003). Multiple-comparisons corrections were applied using an adaptive two-step false discovery rate (FDR; (Benjamini and Hochberg, 2000)), wherein both steps’ maps were thresholded at p<0.01, and the cluster extended spatio-temporal threshold set to 200 voxels and two consecutive time windows. Finally, for visualization purposes, a binarized image generated from patients’ group-level atrophy map was used as an exclusive mask applied to the voxelwise statistics (see section *MRI protocol and analyses*). Figures 2-4 focus on the left hemisphere results, central to our investigation and main component of our results. For completeness, Table 3 reports all statistically significant clusters in both hemispheres.

**Table 3.**
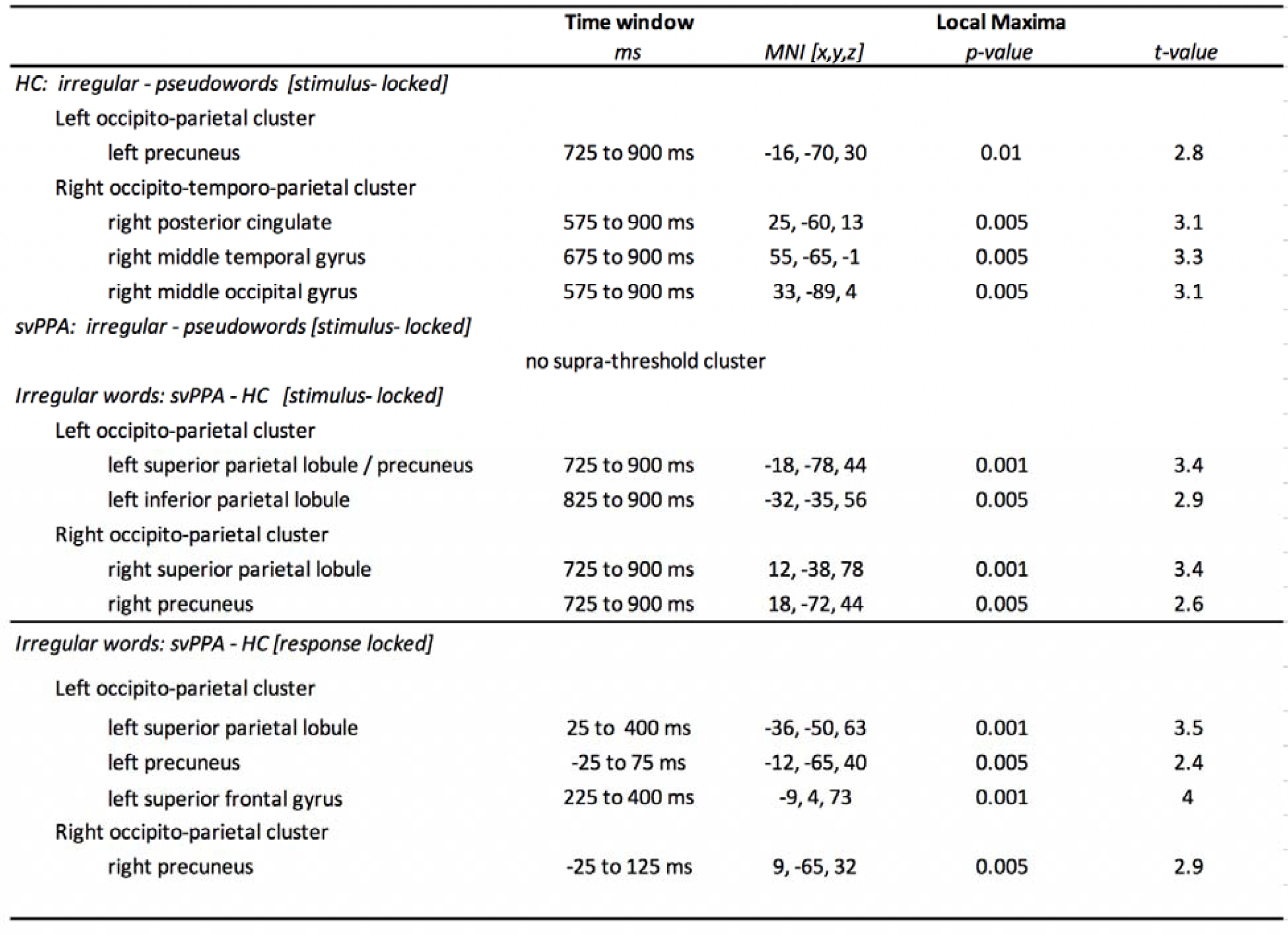
Local maxima in MNI coordinates. Time window, MNI coordinates and t-value of the local maxima of the different MEG whole-brain contrasts performed. The spatiotemporal distribution of these clusters can be appreciated in Fig. 3 and 4.

First, we examined the pattern of activation during irregular word and pseudoword reading separately for controls and svPPA patients (SnPM one-sample t-test against baseline). Then, for both groups, we generated within-subject contrast maps comparing pseudowords and irregular words in order to identify spatiotemporal clusters of heightened activity due **to** sub-lexical processes (SnPM two-sample t-test). Next, we directly compared svPPA patients and controls during irregular word reading in order to highlight spatiotemporal clusters of differential activity between the two groups (SnPM two-sample t-test).

Finally, we conducted two post-hoc analyses. First, we sought to rule out the hypothesis that the differences in neural activation could be explained by differences in reaction times. To this end, the raw datasets were epoched with respect to the onset of vocal response (response-locked trials, from −400 ms before response to +400 ms after vocal response) and the same group comparison (i.e., irregular word reading in controls vs. svPPA patients) was performed. Second, we directly compared, in the time window of interest, svPPA patients and controls during pseudoword reading (SnPM two-sample t-test) to ascertain whether the observed difference between groups was generalized or specific to irregular word reading. This whole-brain analysis was corroborated by a region-of-interest (ROI) follow-up directly contrasting the two cohorts during both irregular and pseudoword reading. Two ROIs were defined, centered on the parietal peak of the contrast svPPA vs. controls during irregular words reading in stimuli-locked data and in response-locked data respectively (stimuli-locked coordinates: [-40 −30], [-40 −30], [55 65], response-locked coordinates: [-40 −30], [-65 −55], [55 65]). Single subjects’ peak beta suppression values during irregular and pseudowords reading were extracted in both ROIs in the 100ms window surrounding the peak effect (stimuli-locked: 725-825 ms, response-locked 25-125 ms). Finally, we explored whole-brain differences between the two cohorts during regular word reading (SnPM two-sample t-test). It should be noted that this contrast is less informative than the previous two given that controls can use both lexico/semantic or sub-lexical/phonological strategies.

### Data Availability

The clinical and neuroimaging data used in the current paper are available from the corresponding author, upon reasonable request. The sensitive nature of patients’ data and our current ethics protocol do not permit open data sharing at this stage.

## Results

### Behavioral data and cortical atrophy

Two ANOVAs based on three word types (regular, irregular, and pseudo-words) and two cohorts (controls vs. svPPA patients) were used to compare behavioral performance in the task during the MEG scan. For overall accuracy (Fig. 1A), there was a significant main effect of cohort (F(1)=29.86, p<0.001), a main effect of word type (F(2)=10.75, p<0.001), as well as a significant interaction (F(2)=3.74, p<0.001). For reaction times (Fig.1B), there was a significant main effect of cohort (F(1)=8.54, p <0.001), a main effect of word type (F(2)=27.82, p=0.0048, but no significant interaction (F(2)=0.12, p =0.88). Post-hoc t-tests directly comparing the two cohorts across the three words types revealed statistically significant differences in accuracy for all conditions [pseudowords (t=2.64, p=0.01), irregular words (t=4.79, p<0.001), regular words (t=2.77, p= 0.01)], and in RTs for irregular words (t = 2.54, p = 0.02). The analyses of error type revealed a significant main effect of cohort (F(1)=102.05, p <0.001), a main effect of word type (F(2)=5.22, p<0.001), and significant interaction between cohort and error type (F(1)=47.76, p<0.001). Post-hoc t-tests highlighted a significant difference between controls and svPPA patients only for regularization errors (t=32.88, p<0.001), and not for lexicalizations (t=1.65, p =0.11). Moreover, the percentage of regularization errors statistically differ from that of lexicalization errors in svPPA patients (t= 9.64, p <0.001). Overall, these results, corroborated by the out-of-scanner neuropsychological data (Table 1), are consistent with a clinical diagnosis of svPPA and a pattern of surface dyslexia.

**Figure 1.**
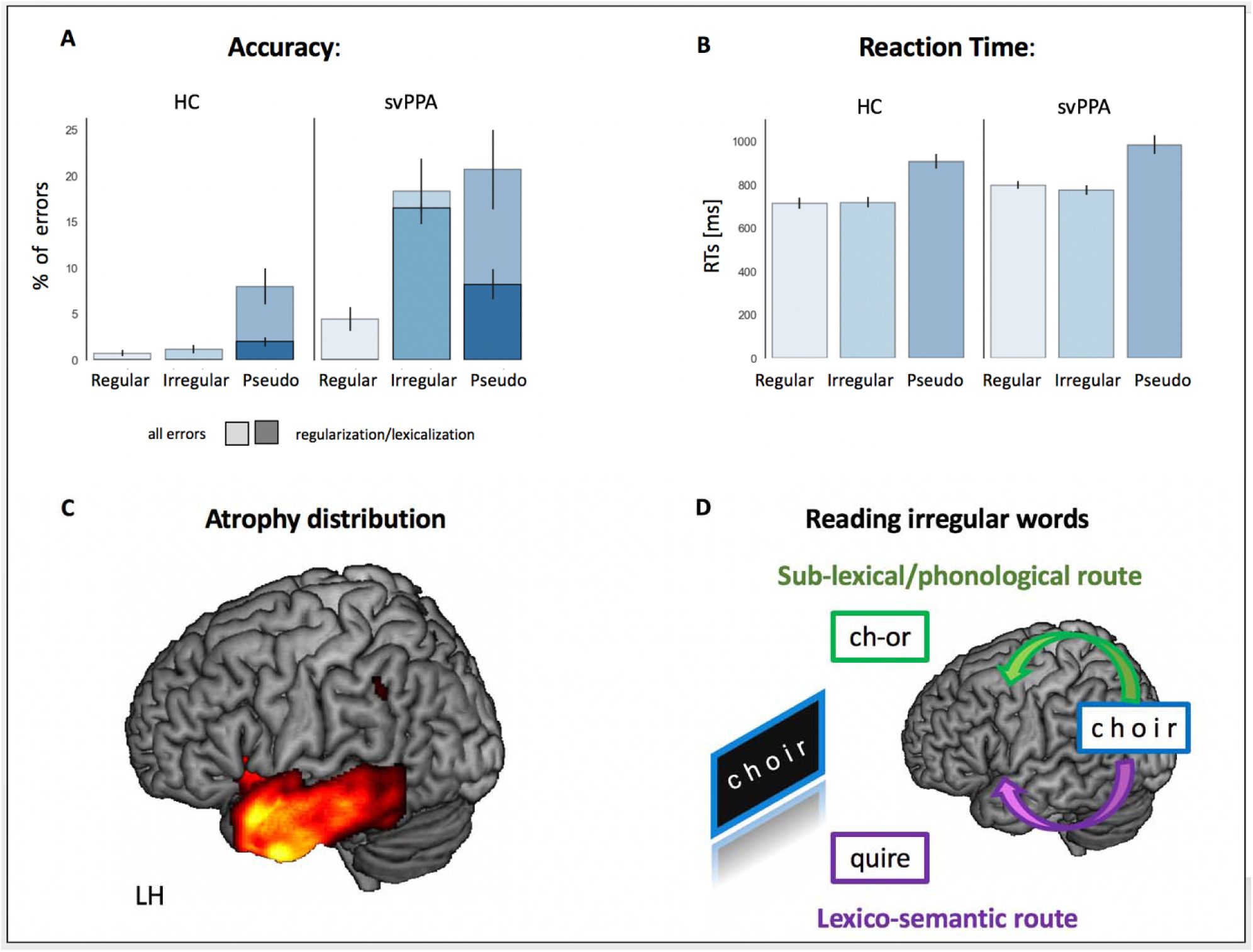
Behavioral data and atrophy distribution in svPPA patients. (A) Percentage of errors in each experimental condition (average and standard error of the mean), for controls (n=12) and svPPA patients (n=11). Darker colors represent the average percentage of regularization errors (with irregular words) and lexicalization errors (in pseudowords). (B) Average reaction times in each experimental condition, in controls (n=12) and svPPA patients (n=11). Errors bars represent the standard error of the mean. (C) Voxel based morphometry (VBM)-derived atrophy pattern showing significantly reduced grey matter volumes in the anterior temporal lobe for svPPA patients (thresholded at p<0.001 with family wise error (FWE) correction). (D) Cartoon representation of the dual route model: the dorsal sub-lexical/phonological route supports orthography-to-phonology mapping based on language-specific statistical rules, while the ventral lexico-semantic route maps orthographic forms to meaning thus allowing correct reading of irregular words.

**Figure 2.**
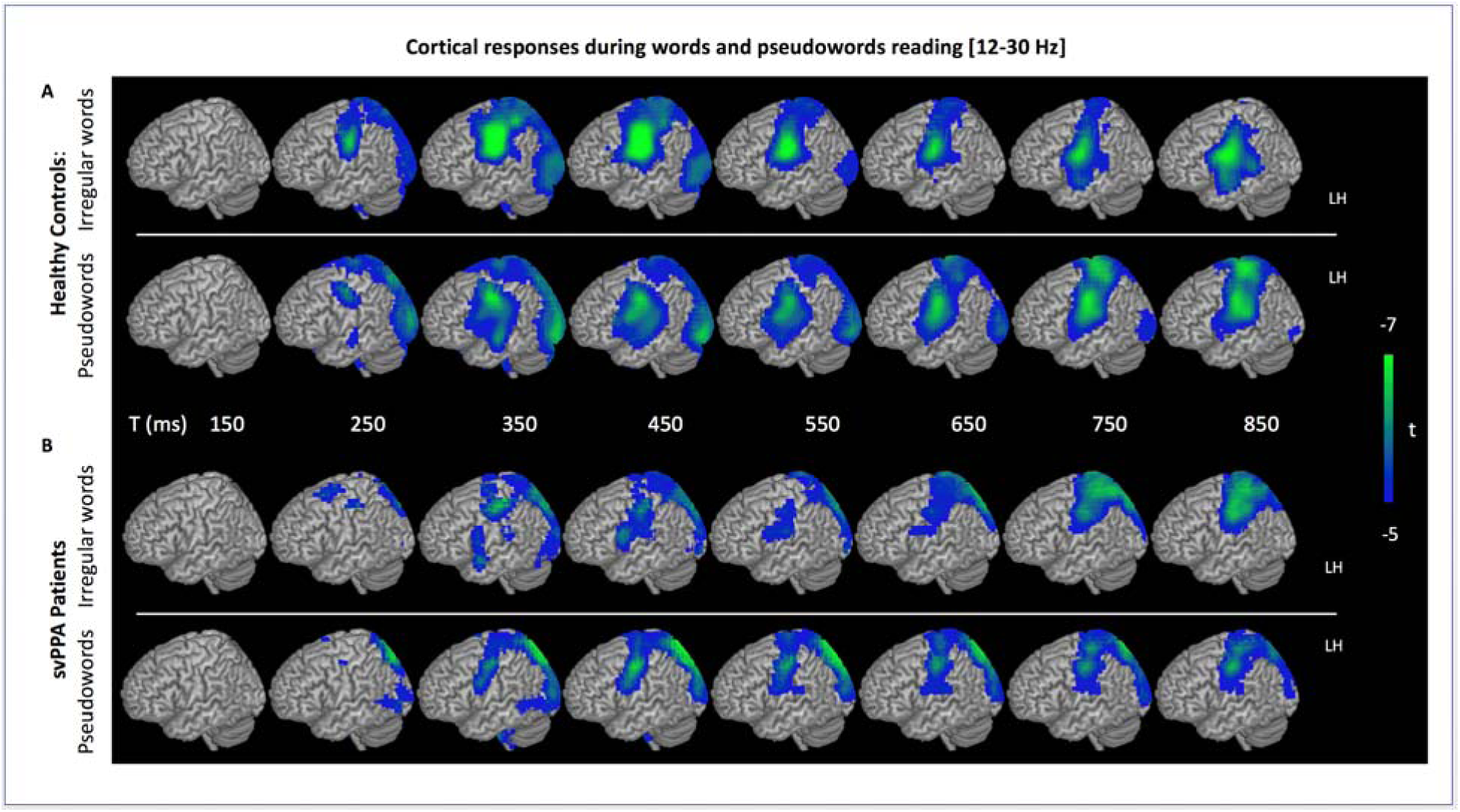
Stimulus-locked (0 ms = written word onset) group analyses of changes in beta (12–30 Hz) oscillatory power during single word reading. A) Pattern of beta suppression in healthy controls for both irregular words (upper row) and pseudowords (lower row) in the left hemisphere. Following stimuli presentation, significantly heightened beta suppression tracks the time course of language network activation from occipital towards temporo-parietal areas. B) Same as in A but for the svPPA patients. Results of one-sample t-test.

Distribution of cortical atrophy in the svPPA cohort is shown in Figure 1C. Patients present the expected pattern of degeneration of the anterior temporal lobe. Unsurprisingly, given our inclusion criteria, the atrophy appears strongly left-lateralized.

### Neural dynamics during reading

Within-group analyses of brain activity (changes in beta power) during irregular word and pseudoword reading are presented for both controls in Figure 2A and svPPA patients in Figure 2B. Following stimuli presentation (0ms), for both groups and conditions we observed significant beta-power suppression (a marker of functional activation) over occipital and parietal areas with a progressive posterior-to-anterior shift in time. This result is coherent with classical findings from both evoked-related fields (Marinkovic *et al.*, 2003) and spectral (Borghesani *et al.*, 2019) analyses.

In order to localize changes in brain activity differential for sub-lexical vs. lexical processes, we then contrasted pseudowords against irregular words separately for each group. In controls, a significant increase in brain activity during pseudoword reading was observed over bilateral occipito-parietal cortices, ramping up ∼600 ms post stimulus onset and peaking at ∼800 ms (Fig 3a). Table 3 summarizes the temporal windows, peaks of local maxima, and t-values of all main sub-clusters that could be isolated using this comparison when corrected for multiple comparisons. This included a cluster over the left precuneus, right posterior cingulate, and right temporal and occipital gyri (Fig 3a). These changes in activation over these parieto-occipital regions peaked during pseudoword reading (when compared to irregular word reading) over these later (600-800ms) time windows, the time of response selection and execution. This finding suggests that, in healthy controls, sub-lexical processes rely on the recruitment of the dorsal route. In svPPA, using the same contrast, this effect was absent even at a very lenient (uncorrected) threshold (Fig 3b). Thus, it appears that, while controls recruit the dorsal route for pseudowords more than for irregular words, svPPA patients process the two types of stimuli in a similar way: relying on the dorsal route.

**Figure 3.**
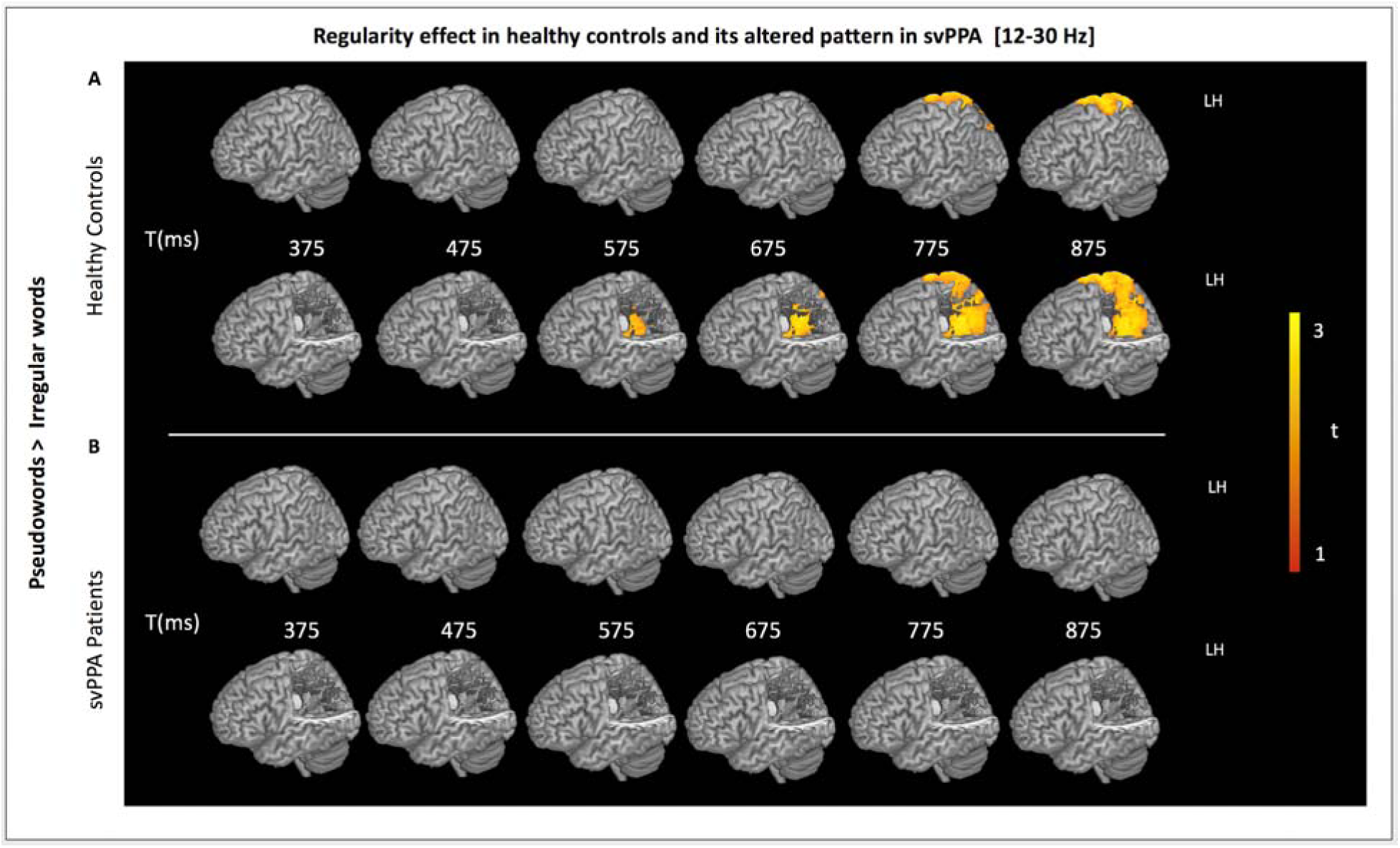
Stimulus-locked (0 ms = written word onset) group analyses of the regularity effect: pseudowords vs. irregular words. A) Rendering of the results of the contrast between pseudowords and irregular words in healthy controls in the left hemisphere with (lower row) and without (upper row) a cut-out allowing appreciation of the effect in the intraparietal sulcus. A large spatiotemporal cluster of heightened beta suppression for pseudowords (vs. irregular words) is observed over the left parietal cortex from ∼ 500ms to ∼ 800 ms. B) Same as in A but for the svPPA patients: no supra-threshold cluster can be detected.

Next, in order to confirm that this increase in activation during sub-lexical processing was absent in svPPA, we directly contrasted irregular word reading in controls against svPPA patients. This direct comparison is crucial as the failure to detect an effect in one of the two groups (i.e., a difference in significance level) does not necessarily entail a significant difference between the two groups (Gelman and Stern, 2006). As we predicted from the separate within-group analyses, this between-group comparison revealed a large spatiotemporal cluster over bilateral occipito-parietal cortex where svPPA patients significantly manifested more brain activity than controls (Fig 4a). As detailed in Table 3, four sub-clusters could be localized, peaking in the bilateral precuneus and superior parietal lobule. The spatial topography and timing of these clusters are highly similar to those described in the within-group analyses, strengthening the interpretation that during irregular word reading, at the time of response selection, patients with svPPA over-recruit the dorsal route. Critically, our post-hoc analysis contrasting the two cohorts in the time windows of interest during pseudoword reading reveal no differences. We corroborated the results of this whole-brain analysis with a region-of-interest follow up, extracting beta suppression values for the two cohorts during both irregular and pseudoword reading. During irregular word reading controls’ average beta suppression was −0.44 (std=0.73), while for patients 1.15 (std=0.75); during pseudoword reading controls’ average was −0.98 (std=0.62), patients’ −0.84 (std=0.91). Statistically, this leads to a significant main effect of cohort (p=0.021), main effect of word type (p=0.006), and of their interaction (p=0.002). Since pseudowords can be read by sub-lexical/phonological route by both cohorts, these findings indicate that patients’ over-recruitment of the dorsal route is specific to irregular word reading and not a generalized reading mechanism. Moreover, ROIs data indicate that none of the participants falls outside of 3 standard deviations for the group mean, thus excluding the possibility that the observed effects are driven by outliers (see Suppl. Fig. 2a). Finally, the comparison svPPA vs. HC during regular word reading isolated a similar spatiotemporal cluster over left parietal cortices as well as a frontal cluster (see Suppl. Fig. 2b). It should be noted that this contrast cannot provide a clear insight on the neural processes underlying lexical/semantic and sub-lexical/phonological processes as these words can be read via either route. However, it still indicates over-reliance on dorsal fronto-parietal cortices as a partial compensation for ventral damage.

**Figure 4.**
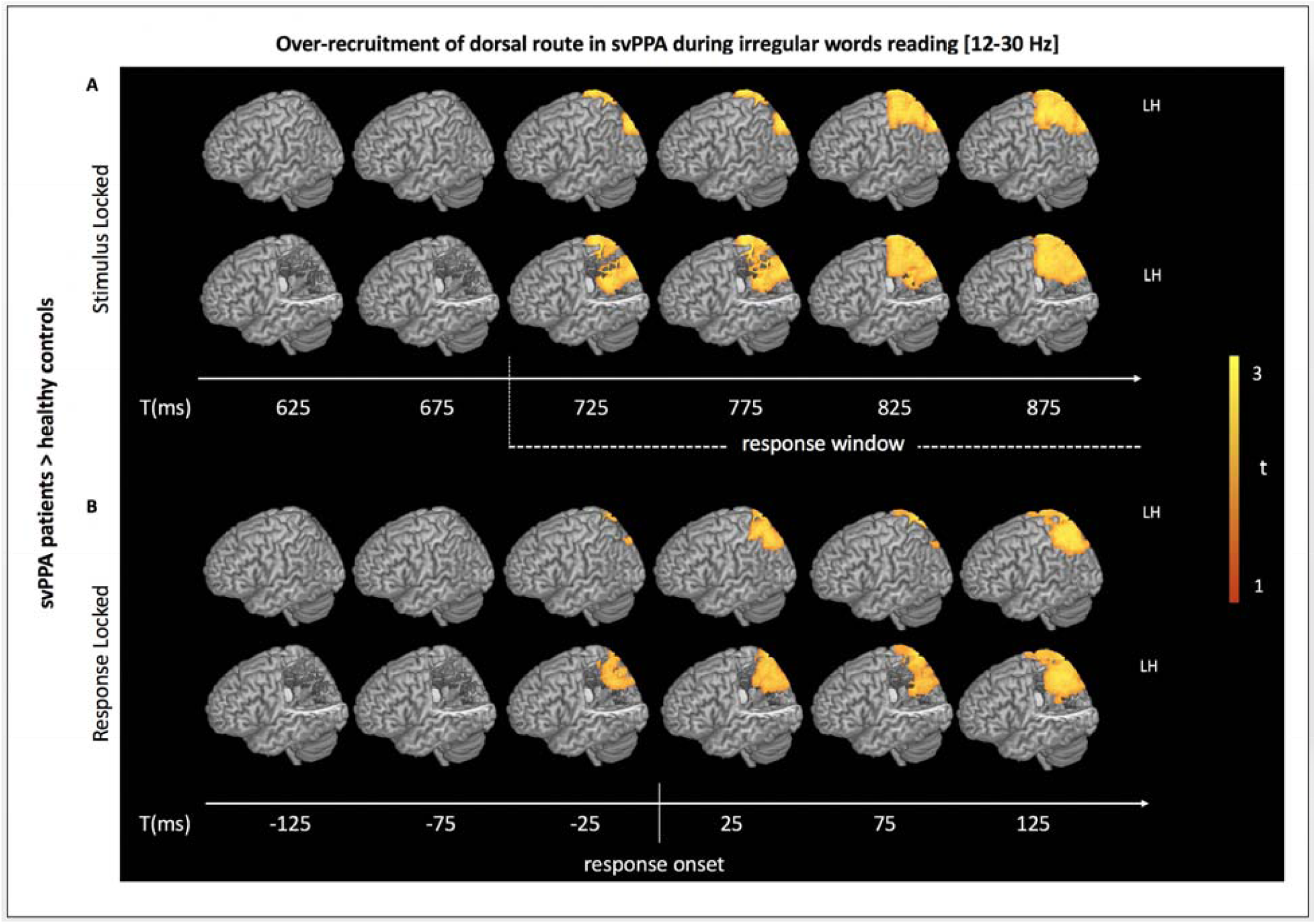
Contrast controls vs. svPPA. A) Rendering of the results of the contrast between svPPA patients and healthy controls during irregular word reading for stimulus-locked analyses (0 ms = written word onset) in the left hemisphere with (lower row) and without (upper row) a cut-out allowing appreciation of the effect in the intraparietal sulcus. A large spatiotemporal cluster of heightened beta suppression in svPPA patients (vs. controls) is observed over the left parietal cortex from ∼ 700ms. B) Same as in A but for response-locked analyses (0 ms = vocal response onset). A similar spatiotemporal cluster is observed starting ∼25 ms before response onset.

To correct for differences in reaction time between the two groups (which could potentially influence our stimulus-locked analyses presented above) we conducted an additional analysis reconstructing changes in brain activity in datasets aligned to the onset of the response (response-locked) when subjects read the word aloud. We conducted a between-group comparison for irregular word reading between controls and svPPA for response-locked trials (Fig 4). Here, we see increases in brain activity similar to the group comparison presented above. Around response onset, significantly increased beta suppression is identifiable in the svPPA group over bilateral occipito-parietal cortices (Fig 4b), in particular involving left precuneus, left superior parietal lobule, and right precuneus (see Table 3 for details). This additional analysis confirms that, even when differences in reaction time are taken into account, patients with svPPA still over-recruit these regions of the parieto-occipital cortex bilaterally.

## Discussion

We provide novel evidence supporting reading models that posit a division of labor between a ventral (lexico-semantic) and a dorsal (sub-lexical/phonological) route based on the spatiotemporal pattern of brain activity associated with reading aloud irregular words and pseudowords in healthy subjects and patients with svPPA. Healthy controls engaged the occipito-parietal cortex to read pseudowords (more than with irregular words), corroborating the involvement of this dorsal route in sub-lexical, orthography-to-phonology processes. In contrast, svPPA patients, who present with surface dyslexia and left anterior temporal atrophy, appeared to recruit this dorsal cluster equally for irregular words and pseudowords. Direct comparison of controls and patients during irregular word reading highlighted the same spatiotemporal cluster. These results support the interpretation that svPPA patients rely on the spared functions underpinned by the dorsal route (i.e., orthography-to-phonology mapping following language-specific statistical regularities) to read in the context of semantic impairment due to ventral route damage. Crucially, these effects were detected at the time of response preparation and production, consistent with the idea that the dorsal sub-lexical route is supporting serial, slow, attention-dependent processes, in contrast with the parallel, fast, automatic ones underpinned by the ventral lexical route.

### Dorsal, occipito-parietal cortices support sub-lexical/phonological processes

This is the first study investigating the spatiotemporal dynamics of reading in svPPA, a clinical syndrome whose neuroanatomical (i.e., neurodegeneration of the ATL) and neuropsychological (i.e., surface dyslexia) features allow scrutiny of the interplay (and dissociation) between lexico-semantic and sub-lexical/phonological processes.

The first main result of our study is that reading pseudowords, as compared with irregular ones, requires heightened recruitment of the parietal cortex. This result is in line with previous neuroimaging and neuropsychological evidence associating the dorsal route with sub-lexical processes (Booth *et al.*, 2003; Jobard *et al.*, 2003; Mechelli *et al.*, 2003; Wilson *et al.*, 2009; Graves *et al.*, 2010; Taylor *et al.*, 2013; Sliwinska *et al.*, 2015). Our results also corroborate previous findings that low-level visual processing of symbolic stimuli in primary and secondary visual areas does not differ between words and orthographically regular pseudowords (Vinckier *et al.*, 2007). At these early stages of visual processing, no differences are detected between healthy controls and svPPA patients, as predicted by the pattern of atrophy and the neuropsychological profile of svPPA patients. However, as compared to healthy controls, svPPA patients do not show heightened recruitment of the parietal cortex when comparing pseudowords and irregular words. This finding suggests that they are processing the two sets of stimuli via the same neuro-cognitive mechanism. This conclusion is supported by the observation that, during irregular word reading, svPPA patients show greater activation (compared to controls) of a very similar parietal cluster. Patients thus appear to rely on dorsal, sub-lexical processes to read both irregular words and pseudowords. Crucially, our observation of a differential recruitment of dorsal cortices in svPPA vs. controls cannot be reduced to an error-related signal since it is observed during irregular words reading but not pseudoword reading. As pseudowords are arguably harder for both groups (i.e., both groups make more errors in pseudowords than words), this finding strengthen the idea that the recruitment of left IPS is modulated by demands on sub-lexical processes rather than a signature of difficulty/errors per se.

The parietal cortex, in particular the inferior parietal lobule, has been associated with various cognitive functions including pointing, grasping, attention orienting, saccades, calculation, and phoneme detection (Simon *et al.*, 2002). With respect to language, discrete areas preferentially involved in orthography and attention, fluency and phonology, or semantics have been identified (Battistella *et al.*, 2019). Overall, bilateral posterior parietal regions appear to support an array of visuo-spatial attentional processes including visual search and scanning of arrays of objects (Corbetta and Shulman, 2002), endogenous orientation of attention to lateralized visual targets (e.g. Gitelman *et al.*, 1999; Peelen *et al.*, 2004), and switching from global (word) to local (letters) attention (Wilkinson *et al.*, 2001). Readers can harness these specific forms of focused attention to apply serial reading strategies if they are young and/or unskilled (Church *et al.*, 2008; Dehaene-Lambertz *et al.*, 2018), or to compensate for visual input degradation (Cohen *et al.*, 2008), brain damage causing word blindness (Cohen *et al.*, 2004; Henry *et al.*, 2005), or loss of lexico-semantic information (Wilson *et al.*, 2009). The role of the parietal cortex as the sub-lexical component of the reading network is further supported by the observation of a correlation between its recruitment and reading proficiency, likely mediated by its structural and functional connectivity profile (Bouhali *et al.*, 2019; Broce *et al.*, 2019; Moulton *et al.*, 2019). Our current investigation lacks appropriate spatial resolution, but future studies deploying other techniques might be able to topographically separate executive/attentional contributions from phonological ones (Oberhuber *et al.*, 2016).

Overall, our findings strengthen the topographical dissociation between (ventral) lexico-semantic and (dorsal) sub-lexical/phonological processes while highlighting the important implications not only for developmental (Pugh *et al.*, 2000; Peyrin *et al.*, 2011) but also acquired reading disorders (Aguilar *et al.*, 2018).

### The latency of the dorsal activation indicates slow, serial processing

The dissociation between lexico-semantic and sub-lexical/phonological processes cannot be resolved in their localization to ventral, occipito-temporal and dorsal, occipito-parietal cortices respectively. Behavioral evidence suggests that the two routes rely on different computational mechanisms and thus have different timing properties (Paap and Noel, 1991; Coltheart and Rastle, 1994; Weekes, 1997). Both routes are thought to be activated by any given written stimulus yet the ventral one would be characterized by fast, parallel processing while the dorsal one by slow, serial processing. Our results, thanks to the temporal resolution offered by MEG recordings, corroborate this perspective.

In healthy controls, the early steps of visual information processing do not differ between irregular words and pseudowords, suggesting that both ventral and dorsal routes are initially equally engaged. Moreover, these early stages do not appear to differ between healthy controls and svPPA patients, consistent with patient’s spared occipital areas and preserved basic visual functions (Watson *et al.*, 2018). Differences between irregular words and pseudowords, as well as between healthy controls and svPPA patients emerge only at the time of response selection and production. In these later, final stages, healthy controls show heightened activity over dorsal areas for sub-lexical/phonological processes (i.e., pseudowords > irregular words). Critically, this regularity effect could not be detected in svPPA patients. Hence, svPPA patients appear to rely on the same neuro-cognitive substrate to process both irregular words and pseudowords. This result is strengthened by the detection of a similar spatiotemporal cluster when directly comparing neural activity during irregular word reading across the two groups. These findings were confirmed by both stimulus-locked and response-locked analyses, indicating that the effects are stable in space and time, and cannot be ascribed to differences in reaction time. Thus, in svPPA, surface dyslexia appears to be due to the failure of activating semantic representations stored in the temporal lobe, and the need to rely on sub-lexical grapheme-to-phoneme translation supported by the parietal lobe.

Our findings are in line with converging behavioral and neuroimaging evidence supporting the idea that, while activated at the same time by the presentation of a written stimulus, the lexical (ventral) route and the sub-lexical (dorsal) one are characterized by different processing dynamics: fast and parallel in the first case, slow and serial in the latter. For instance, the systematic length effect detected for pseudowords is stronger than for real words (Weekes, 1997). Moreover, the strength of the regularity effect declines as a function of the position of the irregular grapheme: reaction times are longer when reading “chef” (i.e., irregular grapheme in first position), than “tomb” (i.e., irregular grapheme in second position), than “glow” (i.e., irregular grapheme in third position) (Coltheart and Rastle, 1994). Electroencephalography (EEG), magnetoencephalography (MEG), and intracranial local field potentials (LFP) have been instrumental in illustrating how basic visual feature analysis occurs within ∼100 ms over the occipital midline, followed by a word-selective yet pre-lexical response peaking over the left occipital scalp around 150-200 ms (Bentin *et al.*, 1999; Tarkiainen *et al.*, 1999; Gaillard *et al.*, 2006). These initial sequential stages are followed by parallel processing of nonvisual information in a distributed language network allowing access to lexical, semantic and phonological features of the words (Vinckier *et al.*, 2007; Twomey *et al.*, 2011; Thesen *et al.*, 2012). On one hand, semantic effects are typically detected at around 400-600 ms (e.g. N400) and localized to left ATL (Lau *et al.*, 2008, 2016). On the other hand, consistent with the role played by the parietal lobe in visual attention (Saalmann *et al.*, 2007), late sustained engagement of dorsal regions is observed in cases of effortful reading (Rosazza *et al.*, 2009).

The effects we observe occur close to response production and are strongest after response onset - i.e., when the subjects are spelling out the word. The progressively heightened activity over dorsal cortices suggests that, when relying on the sub-lexical route, articulation of the word starts as the first grapheme-to-phoneme conversion has been performed and proceeds serially grapheme by grapheme. This is in contrast with the parallel processing taking place along the ventral lexical route that allows spoken production once the stimulus has been decoded as a whole. Overall, our findings corroborate the hypothesis that the dorsal sub-lexical route supports serial and attention-dependent processes, whose effects are most apparent at the time of response selection and production.

### Over-recruitment of the dorsal route as the neural signatures of an imperfect compensation

Our results illustrate how a specific cognitive task can be performed via the deployment of distinct cognitive and neural resources. The damage to the ATL in svPPA patients leads to a profound impairment of the lexico-semantic system, whose functionality is needed to retrieve the correct pronunciation of exceptional spelling-to-sound associations. As a compensatory strategy, svPPA patients rely on sub-lexical processes underpinned by the dorsal route to read irregular words. However, patients’ over-reliance on serial, slow, grapheme-to-phoneme conversion mechanisms leads to overall slower responses and, in the case of words with atypical spelling, increased number of regularization errors. Results from this study demonstrate that patients can thus only partially, imperfectly compensate for their semantic loss via spared sub-lexical processes.

Our findings align with recent evidence from task-free fMRI studies reporting alterations in intrinsic functional connectivity networks in svPPA patients. Decreased connectivity in the ventral semantic network (as expected given the atrophy pattern) is accompanied by spared connectivity in the orthography-to-phonology conversion network and increased connectivity in the dorsal articulatory-phonological one (Battistella *et al.*, 2019; Montembeault *et al.*, 2019). Taken together, these results suggest that neurodegeneration leads to a dynamic reorganization of the interplay between ventral and dorsal pathways, where the downregulation of specific neuro-cognitive systems is associated with the upregulation of other ones. Moreover, they exemplify how studying the spatiotemporal dynamics of different language functions in clinical populations has implications for both clinical practice (e.g., understanding patients’ symptoms and informing rehabilitation approaches) and basic science (e.g., understanding the neural correlates of language). These studies also stress the need to shift the focus of research in neurodegenerative populations from volume and structural connection loss to also include alterations to regions’ functional profiles and temporal dynamics.

MEG has only been deployed recently to investigate PPA (e.g., Kielar *et al.*, 2018), but has already shown its potential as a tool to study the neurophysiological signatures of network-level alterations due to neurodegenerative disorders. It can be instrumental in identifying syndrome-specific changes in the spectral properties of oscillatory responses (Ranasinghe *et al.*, 2017; Sami *et al.*, 2018). As suggested by task-free fMRI evidence, these functional alterations might precede structural ones and may be key for early diagnosis (Bonakdarpour *et al.*, 2017). Given the different features of the signal that can be studied (e.g., delayed latencies, attenuated amplitude) and the wide array of analysis methods available, MEG will provide a broad spectrum of complementary insights into the neural underpinnings of neurodegenerative diseases.

Patients with stroke affecting the left hemisphere might exhibit a different neuropsychological profile from svPPA ones: regularization errors do not appear to correlate with semantic impairments and are associated with damage to the posterior half of the left middle temporal gyrus rather than the ATL (Binder *et al.*, 2016). This finding suggests that the neuro-cognitive profile of svPPA patients (i.e., surface dyslexia, semantic deficits, and ATL damage) is not the only path to over-reliance on sub-lexical processes. Additional functional studies comparing stroke and neurodegenerative patients are warranted to clarify whether the spatio-temporal dynamics of the compensatory mechanisms implemented by the two clinical populations are similar.

### Limitations and future perspectives

Our sample size is relatively small. However, svPPA is a rare disorder and only a fraction of patients can surmount the difficulties inherent to MEG testing (e.g., percentage of dental work, motion, fatigue). However, only a larger patient sample will allow future studies to investigate the relation between specific error types and regional activity. In the present study, our capacity to contrast predictions from different cognitive accounts of reading (e.g., DRC vs. triangle model) is constrained not only by our statistical power, but also by our stimuli choice. For instance, we cannot disentangle the contribution of regularity from that of frequency and consistency (see Suppl. Fig.1). Finally, we optimized our analysis pipeline to investigate possible late effects of the key contrast of interest (irregular vs pseudowords) as prior findings suggested a key role of dorsal, sub-lexical mechanisms in svPPA patients with surface dyslexia (Wilson *et al.*, 2009). Follow-up studies of other electrophysiological features (e.g., lower frequencies, evoked fields) might reveal earlier, additional effects.

### Conclusions

Together these findings, combining a neuropsychological model and time-resolved MEG imaging, provide further evidence supporting a dual-route model for reading aloud, mediated by the interplay between lexical (ventral and parallel) and sub-lexical (dorsal and serial) neuro-cognitive systems. When the lexical route is damaged, as in the case of neurodegeneration affecting the ATL, partial compensation appears to be possible by over-recruitment of the slower, serial attention-dependent, sub-lexical one. This study also demonstrates how task-based MEG imaging can be leveraged in clinical populations to study the functional interplay of different language networks, with important implications for both models of language and clinical practice.

## Acknowledgements

The authors thank the patients and their families for the time and effort they dedicated to this research.

---

**Suppl. Fig. 1.**
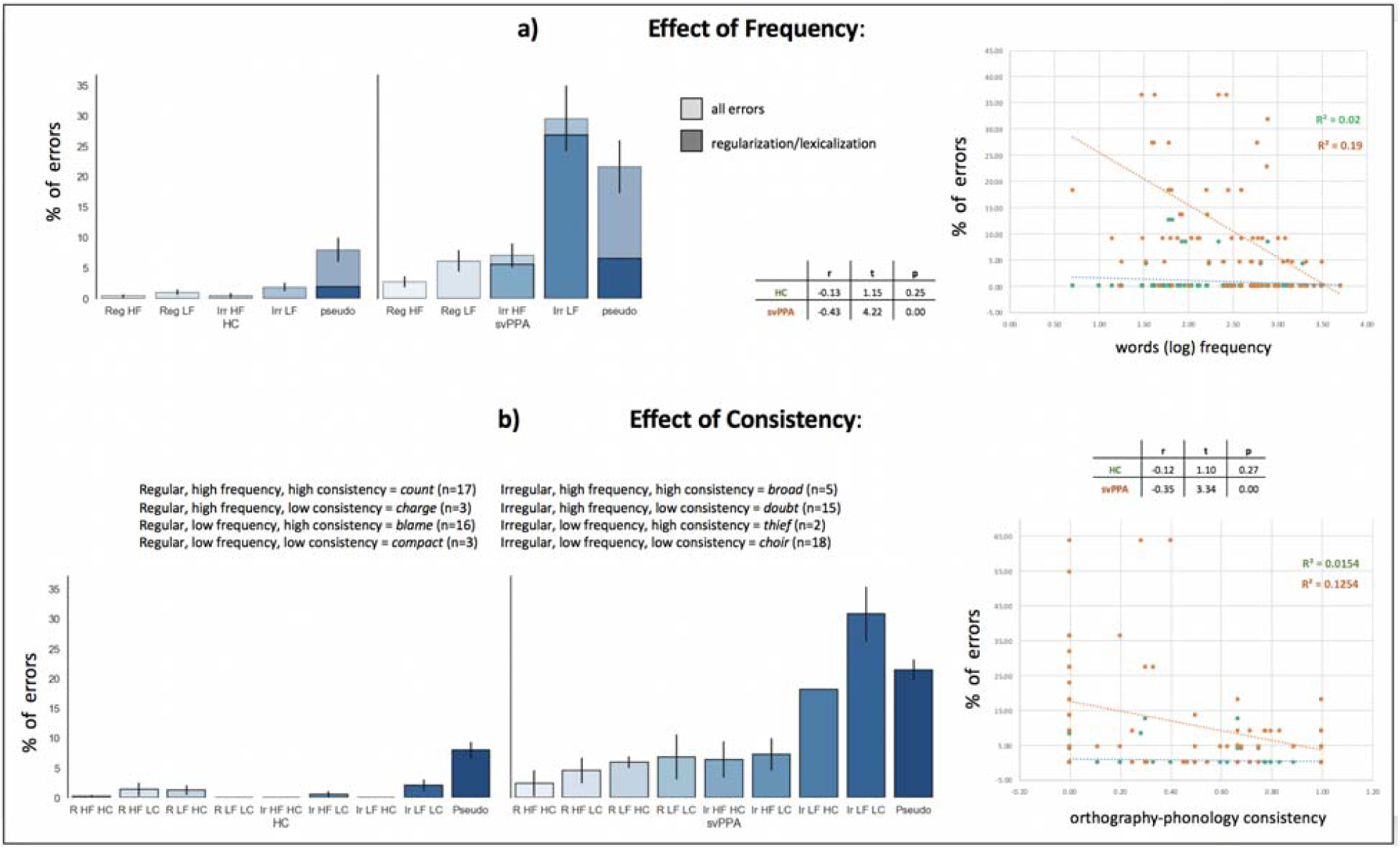
Analysis of the effect of words frequency and orthographic-to-phonological consistency. (A) On the left, percentage of errors (average and standard error of the mean), for controls (n=12) and svPPA patients (n=11) across 5 conditions: regular high frequency words, regular low frequency words, irregular high frequency words, irregular low frequency words, pseudowords. Darker colors represent the average percentage of regularization errors (with irregular words) and lexicalization errors (in pseudowords). An ANOVA 2 (cohort) * 5 (word type) revealed a significant main effect of cohort (F(1)=49.92, p<0.001), a main effect of word type (F(4)=14.20, p<0.001), as well as a significant interaction (F(4)=8.43, p <0.001). On the right, scatterplot illustrating the linear relation between word frequency and percentage of errors in the two cohorts. The correlation is significant in svPPA patients (r = −0.43, p <0.001). (B) On the left, percentage of errors (average and standard error of the mean), for controls (n=12) and svPPA patients (n=11) across 9 conditions: regular high frequency high consistency words, regular high frequency low consistency words, regular low frequency high consistency words, regular low frequency low consistency words, irregular high frequency high consistency words, irregular high frequency low consistency words, irregular low frequency high consistency words, irregular low frequency low consistency words, pseudowords. On the right, scatterplot illustrating the linear relation between word orthography-to-phonology consistency and percentage of errors in the two cohorts. The correlation is significant in svPPA patients (r = - 0.35, p <0.001).

**Suppl. Fig. 2.**
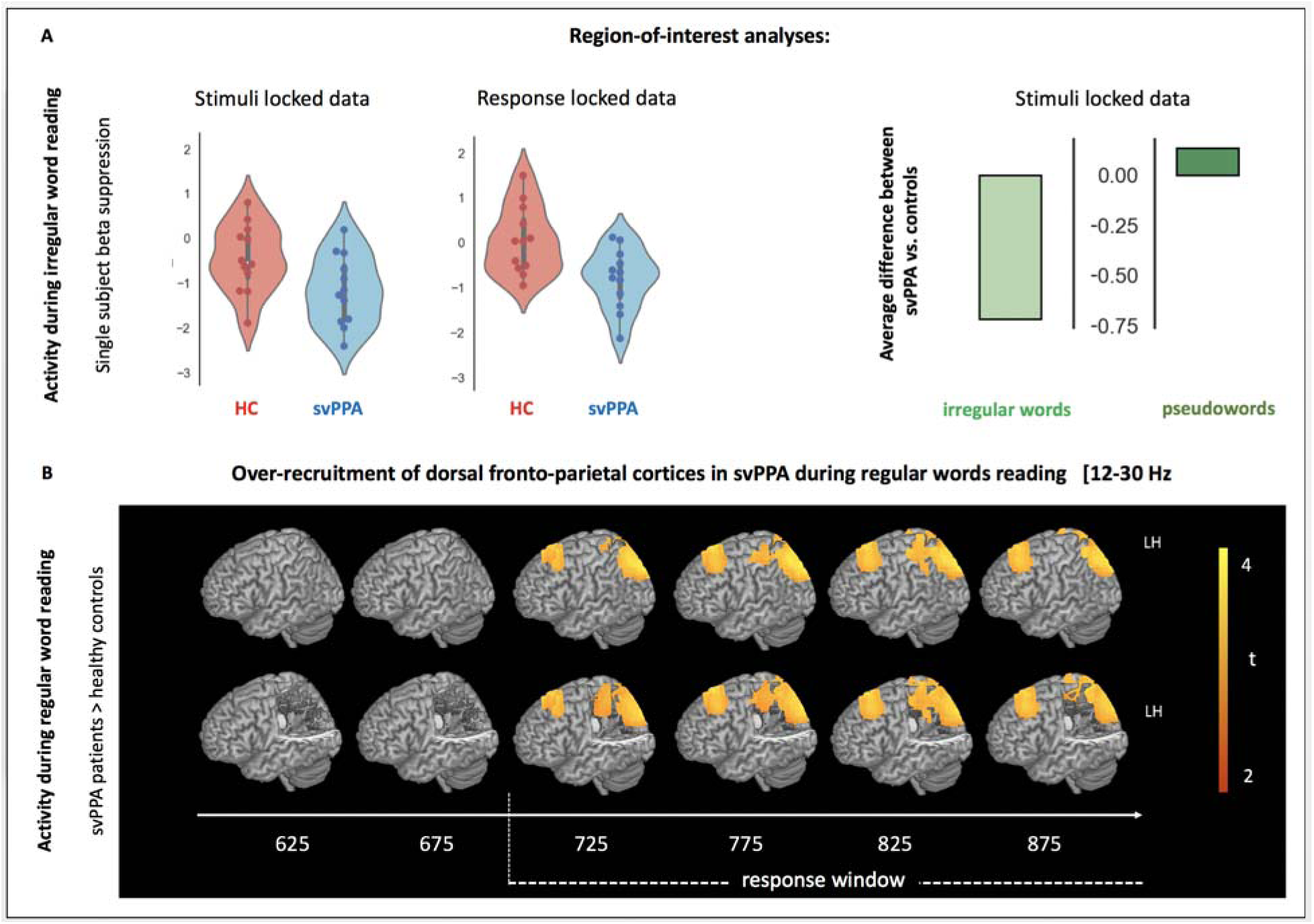
Analyses corroborating the main MEG effects. A) Region-of-interest analyses. Left: single subjects’ beta suppression values during irregular word reading extracted from the parietal peak of the group contrast (stimuli-locked data coordinates: [-40 −30], [-40 −30], [55 65], and time window: 725-825 ms; response-locked data coordinates: [-40 −30], [-65 −55], [55 65]) and time window: 25-125 ms). None of the participants falls outside of 3 standard deviations for the group mean, thus excluding the possibility that the observed effects are driven by outliers. Moreover, a significant main effect of cohort (p=0.021), main effect of word type (p=0.006), and of their interaction (p=0.002) indicate that patients’ over-recruitment of the dorsal route is specific to irregular word reading and not a generalized reading mechanism. Right: graphical illustration of the difference between controls and svPPA patients during irregular word and pseudoword reading: the two cohorts statistically differ only during irregular word reading (when svPPA patients have significantly higher values of beta suppressions, see text for details). B) Rendering of the results of the contrast between svPPA patients and healthy controls during regular word reading for stimulus-locked analyses (0 ms = written word onset) in the left hemisphere with (lower row) and without (upper row) a cut-out. Compared to controls, svPPA patients present large spatiotemporal clusters of heightened beta suppression over fronto-parietal, dorsal cortices. The timing and the location of the parietal cluster overlap with the one isolated by the contrast svPPA vs. controls during irregular word reading (Figure 4).

## References

Aguilar OM, Kerry SJ, Crinion JT, Callaghan MF, Woodhead ZVJ, Leff AP. Dorsal and ventral visual stream contributions to preserved reading ability in patients with central alexia. Cortex 2018; 106: 200–212.

Balota, David A., Melvin J. Yap, Keith A. Hutchison, Michael J. Cortese, Brett Kessler, Bjorn Loftis, James H. Neely, Douglas L. Nelson, Greg B. Simpson, and Rebecca Treiman. “The English lexicon project.” Behavior research methods 39, no. 3 (2007): 445–459.

Battistella G, Henry M, Gesierich B, Wilson SM, Borghesani V, Shwe W, et al. Differential intrinsic functional connectivity changes in semantic variant primary progressive aphasia. NeuroImage Clin 2019; 22: 101797.

Beeson PM, Rising K. Arizona battery for reading and spelling. 2010

Benjamini Y, Hochberg Y. On the Adaptive Control of the False Discovery Rate in Multiple Testing With Independent Statistics. J Educ Behav Stat 2000; 25: 60–83.

Bentin S, Mouchetant-Rostaing Y, Giard M, Echallier J, Pernier J. ERP manifestations of processing printed words at different psycholinguistic levels: time course and scalp distribution. J Cogn Neurosci 1999; 11: 235–260.

Binder JR, Pillay SB, Humphries CJ, Gross WL, Graves WW, Book DS. Surface errors without semantic impairment in acquired dyslexia: A voxel-based lesion-symptom mapping study. Brain 2016; 139: 1517–1526.

Binney RJ, Henry ML, Babiak M, Pressman PS, Santos-Santos MA, Narvid J, et al. Reading words and other people: A comparison of exception word, familiar face and affect processing in the left and right temporal variants of primary progressive aphasia. Cortex 2016; 82: 147–163.

Bonakdarpour B, Rogalski EJ, Wang A, Sridhar J, Mesulam MM, Hurley RS. Functional Connectivity is Reduced in Early-stage Primary Progressive Aphasia When Atrophy is not Prominent. Alzheimer Dis Assoc Disord 2017; 31: 101–106.

Booth JR, Burman DD, Meyer JR, Gitelman DR, Parrish TB, Mesulam MM. Relation between brain activation and lexical performance. Hum Brain Mapp 2003; 19: 155–169.

Borghesani V, Buiatti M, Eger E, Piazza M. Conceptual and Perceptual Dimensions of Word Meaning Are Recovered Rapidly and in Parallel during Reading. J Cogn Neurosci 2019; 26: 194–198.

Bouhali F, Bézagu Z, Dehaene S, Cohen L. A mesial-to-lateral dissociation for orthographic processing in the visual cortex. PNAS 2019; 116: 21936–21946.

Brambati SM, Ogar J, Neuaus J, Miller B, Gorno-Tempini M. Reading Disorders in Primary Progressive Aphasia: a behavioral and neuroimaging study. Neuropsychologia 2009; 47: 1893–1900.

Brambati SM, Rankin KP, Narvid J, Seeley WW, Dean D, Rosen HJ, et al. Atrophy progression in semantic dementia with asymmetric temporal involvement: A tensor-based morphometry study. Neurobiol Aging 2009; 30: 103–111.

Broce IJ, Bernal B, Altman N, Bradley C, Baez N, Cabrera L, et al. Fiber pathways supporting early literacy development in 5–8-year-old children. Brain Cogn 2019; 134: 80–89.

Cai C, Xu J, Velmurugan J, Knowlton R, Sekihara K, Nagarajan SS, et al. Evaluation of a dual signal subspace projection algorithm in magnetoencephalographic recordings from patients with intractable epilepsy and vagus nerve stimulators. Neuroimage 2018; 188: 161–170.

Church JA, Coalson RS, Lugar HM, Petersen SE, Schlaggar BL. A developmental fMRI study of reading and repetition reveals changes in phonological and visual mechanisms over age. Cereb Cortex 2008; 18: 2054–2065.

Cohen L, Dehaene S, Vinckier F, Jobert A, Montavont A. Reading normal and degraded words: Contribution of the dorsal and ventral visual pathways. Neuroimage 2008; 40: 353–366.

Cohen L, Henry C, Dehaene S, Martinaud O, Lehéricy S, Lemer C, et al. The pathophysiology of letter-by-letter reading. Neuropsychologia 2004; 42: 1768–1780.

Coltheart M. Acquired dyslexias and the computational modelling of reading. Cogn Neuropsychol 2006; 23: 96–109.

Coltheart M, Curtis B, Atkins P, Haller M. Models of reading aloud. Psychol Rev 1993; 100: 589–608.

Coltheart M, Rastle K. Serial Processing in Reading Aloud: Evidence for Dual-Route Models of Reading. J Exp Psychol Hum Percept Perform 1994; 20: 1197–1211.

Coltheart M, Rastle K, Perry C, Langdon R, Ziegler J. DRC: a dual route cascaded model of visual word recognition and reading aloud. Psychol Rev 2001; 108: 204.

Corbetta M, Shulman GL. Control of goal-directed and stimulus-driven attention in the brain. Nat Rev Neurosci 2002; 3: 201–215.

Dalal SS, Guggisberg AG, Edwards E, Sekihara K, Findlay AM, Canolty RT, et al. Five-dimensional neuroimaging: Localization of the time-frequency dynamics of cortical activity. Neuroimage 2008; 40: 1686–1700.

Dehaene-Lambertz G, Monzalvo K, Dehaene S. The emergence of the visual word form: Longitudinal evolution of category-specific ventral visual areas during reading acquisition. 2018

Gaillard R, Naccache L, Pinel P, Clémenceau S, Volle E, Hasboun D, et al. Direct Intracranial, fMRI, and Lesion Evidence for the Causal Role of Left Inferotemporal Cortex in Reading. Neuron 2006; 50: 191–204.

Galton C, Patterson K, Graham K, Lambon-Ralph M, Williams G, Antoun N, et al. Differing patterns of temporal atrophy in Alzheimer’s disease and semantic Differing patterns of temporal atrophy in Alzheimer’s disease and semantic dementia. Neurology 2001; 57: 216–225.

Gelman A, Stern H. The difference between ‘significant’ and ‘not significant’ is not itself statistically significant. Am Stat 2006; 60: 328–331.

Gitelman DR, Nobre AC, Parrish TB, LaBar KS, Kim YH, Meyer JR, et al. A large-scale distributed network for covert spatial attention. Further anatomical delineation based on stringent behavioural and cognitive controls. Brain 1999; 122: 1093–1106.

Gorno-Tempini M, Dronkers N, Rankin K, Ogar J, Phengrasamy L, Rosen H, et al. Cognition and anatomy in three variants of primary progressive aphasia. Ann Neurol 2004; 55: 335–346.

Gorno-tempini ML, Hillis AE, Weintraub S, Kertesz A, Mendez M, Cappa S, et al. Classification of primary progressive aphasia and its variants. Neurology 2011; 02: 1–10.

Graham NL, Patterson K, Hodges JR. The impact of semantic memory impairment on spelling: Evidence from semantic dementia. Neuropsychologia 2000; 38: 143–163.

Grainger J, Holcomb PJ. Watching the Word Go by: On the Time-course of Component Processes in Visual Word Recognition. Lang Linguist Compass 2009; 3: 128–156.

Graves WW, Desai R, Humphries C, Seidenberg MS, Binder JR. Neural systems for reading aloud: A multiparametric approach. Cereb Cortex 2010; 20: 1799–1815.

Henry C, Gaillard R, Volle E, Chiras J, Ferrieux S, Dehaene S, et al. Brain activations during letter-by-letter reading: A follow-up study. Neuropsychologia 2005; 43: 1983–1989.

Herman AB, Houde JF, Vinogradov S, Nagarajan SS. Parsing the phonological loop: Activation timing in the dorsal speech stream determines accuracy in speech reproduction. J Neurosci 2013; 33: 5439–5453.

Hinkley LBN, Marco EJ, Brown EG, Bukshpun P, Gold J, Hill S, et al. The Contribution of the Corpus Callosum to Language Lateralization. J Neurosci 2016; 36: 4522–4533.

Hodges J, Garrard P, Perry R, Patterson K, Bak T, Gregory C. The differentiation of semantic dementia and frontal lobe dementia from early Alzheimer’s disease: a comparative neuropsychological study. Neuropsychology 1999; 13: 31–40.

Hodges JR, Patterson K, Oxbury S, Funnell E. Semantic dementia. Progressive fluent aphasia with temporal lobe atrophy. Brain 1992; 115: 1783–1806.

Hoffman P, Lambon Ralph MA, Woollams AM. Triangulation of the neurocomputational architecture underpinning reading aloud. Proc Natl Acad Sci 2015; 112: E3719–E3728.

Jefferies E, Lambon-Ralph MA, Jones R, Bateman D, Patterson K. Surface dyslexia in semantic dementia: A comparison of the influence of consistency and regularity. Neurocase 2004; 10: 290–299.

Jobard G, Crivello F, Tzourio-Mazoyer N. Evaluation of the dual route theory of reading: A metanalysis of 35 neuroimaging studies. Neuroimage 2003; 20: 693–712.

Keuleers E, Brysbaert M. Wuggy: A multilingual pseudoword generator. Behav Res Methods 2010; 42: 627–633.

Kielar A, Deschamps T, Jokel R, Meltzer JA. Abnormal language-related oscillatory responses in primary progressive aphasia. NeuroImage Clin 2018; 18: 560–574.

Kramer JH, Jurik J, Sha SJ, Rankin KP, Rosen HJ, Johnson JK, et al. Distinctive Neuropsychological Patterns in Frontotemporal Dementia, Semantic Dementia, and Alzheimer Disease. Cogn Behav Neurol 2003; 16: 211–218.

Kucera, H., & Francis, W. (1967). Computational analysis of presentday American English. Providence, RI: Brown University Press.

Lau E, Phillips C, Poeppel D. A cortical network for semantics: (de)constructing the N400. Nat Rev Neurosci 2008; 9: 920–933.

Lau E, Weber K, Gramfort A, Hämäläinen M, Kuperberg G. Spatiotemporal Signatures of Lexical-Semantic Prediction. Cereb Cortex 2016; 26: 1377–1387.

Lund, K., & Burgess, C. (1996) Producing high-dimensional semantic spaces from lexical co-occurrence. Behavior Research Methods, Instruments, & Computers, 28, 203–208.

Marelli, M., Amenta, S. A database of orthography-semantics consistency (OSC) estimates for 15,017 English words. Behav Res 50, 1482–1495 (2018). https://doi.org/10.3758/s13428-018-1017-8

Marinkovic K, Dhond RP, Dale AM, Glessner M, Carr V, Halgren E. Spatiotemporal dynamics of modality-specific and supramodal word processing. Neuron 2003; 38: 487–497.

Marshall JC, Newcombe F. Patterns of paralexia: A psycholinguistic approach. J Psycholinguist Res 1973; 2: 175–199.

Mechelli A, Gorno-Tempini ML, Price CJ. Neuroimaging Studies of Word and Pseudoword Reading: Consistencies, Inconsistencies, and Limitations. J Cogn Neurosci 2003; 15: 260–271.

Montembeault M, Chapleau M, Jarret J, Boukadi M, Laforce R, Wilson MA, et al. Differential language network functional connectivity alterations in Alzheimer’s disease and the semantic variant of primary progressive aphasia. Cortex 2019; 117: 284–298.

Moulton E, Bouhali F, Monzalvo K, Poupon C, Zhang H, Dehaene S, et al. Connectivity between the visual word form area and the parietal lobe improves after the first year of reading instruction□: a longitudinal MRI study in children. Brain Struct Funct 2019; 224: 1519–1536.

Neary D, Snowden JS, Gustafson L, Passant U, Stuss D, Black S, et al. Frontotemporal lobar degeneration: A consensus on clinical diagnostic criteria. Neurology 1998; 51: 1546–1554.

Oberhuber M, Hope TMH, Seghier ML, Parker Jones O, Prejawa S, Green DW, et al. Four Functionally Distinct Regions in the Left Supramarginal Gyrus Support Word Processing. Cereb Cortex 2016; 26: 4212–4226.

Paap KR, Noel RW. Dual-route models of print to sound: Still a good horse race. Psychol Res 1991; 53: 13–24.

Patterson K, Hodges JR. Deterioration of word meaning: Implications for reading. Neuropsychologia 1992; 30: 1025–1040.

Patterson K, Ralph MAL, Jefferies E, Woollams A, Jones R, Hodges JR, et al. ‘Presemantic’ cognition in semantic dementia: Six deficits in search of an explanation. J Cogn Neurosci 2006; 18: 169–183.

Peelen M V., Heslenfeld DJ, Theeuwes J. Endogenous and exogenous attention shifts are mediated by the same large-scale neural network. Neuroimage 2004; 22: 822–830.

Perry C, Ziegler JC, Zorzi M. Nested incremental modeling in the development of computational theories: the CDP+ model of reading aloud. Psychol Rev 2007; 114: 273–315.

Peyrin C, Démonet JF, N’Guyen-Morel MA, Le Bas JF, Valdois S. Superior parietal lobule dysfunction in a homogeneous group of dyslexic children with a visual attention span disorder. Brain Lang 2011; 118: 128–138.

Piai V, Roelofs A, Rommers J, Maris E. Beta oscillations reflect memory and motor aspects of spoken word production. Hum Brain Mapp 2015; 36: 2767–2780.

Plaut, D. C., McClelland, J. L., Seidenberg, M. S., & Patterson, K. (1996). Understanding normal and impaired word reading: computational principles in quasi-regular domains. Psychological review, 103(1), 56.

Provost JS, Brambati SM, Chapleau M, Wilson MA. The effect of aging on the brain network for exception word reading. Cortex 2016; 84: 90–100.

Pugh KR, Mencl WE, Jenner AR, Katz L, Frost SJ, Lee JR, et al. Functional neuroimaging studies of reading and reading disability (developmental dyslexia). Ment Retard Dev Disabil Res Rev 2000; 6: 207–213.

Ranasinghe KG, Hinkley LB, Beagle AJ, Mizuiri D, Honma SM, Welch AE, et al. Distinct spatiotemporal patterns of neuronal functional connectivity in primary progressive aphasia variants. Brain 2017; 140: 2737–2751.

Robinson S, Vrba J. Functional neuroimaging by synthetic aperture magnetometry (SAM). In: Yoshimoto, editor(s). Recent advances in biomagnetism. Sendai, Japan: Tohoku University; 1999. p. 302–305

Rosazza C, Cai Q, Minati L, Paulignan Y, Nazir TA. Early involvement of dorsal and ventral pathways in visual word recognition: An ERP study. Brain Res 2009; 1272: 32–44.

Rosen HJ, Gorno-Tempini ML, Goldman WP, Perry RJ, Schuff N, Weiner M, et al. Patterns of brain atrophy in frontotemporal dementia and semantic dementia. Neurology 2002; 58: 198–208.

Saalmann YB, Pigarev IN, Vidyasagar TR. Neural Mechanisms of Visual Attention: How Top-Down Feedback. Science (80-) 2007: 1612–1616.

Salmelin R. Clinical neurophysiology of language: The MEG approach. Clin Neurophysiol 2007; 118: 237–254.

Sami S, Williams N, Hughes LE, Cope TE, Rittman T, Coyle-Gilchrist ITS, et al. Neurophysiological signatures of Alzheimer’s disease and frontotemporal lobar degeneration: Pathology versus phenotype. Brain 2018; 141: 2500–2510.

Sekihara K, Kawabata Y, Ushio S, Sumiya S, Kawabata S, Adachi Y, et al. Dual signal subspace projection (DSSP): A novel algorithm for removing large interference in biomagnetic measurements. J Neural Eng 2016; 13

Simon O, Mangin JF, Cohen L, Le Bihan D, Dehaene S. Topographical layout of hand, eye, calculation, and language-related areas in the human parietal lobe. Neuron 2002; 33: 475–487.

Singh KD, Barnes GR, Hillebrand A. Group imaging of task-related changes in cortical synchronisation using nonparametric permutation testing. Neuroimage 2003; 19: 1589–1601.

Sliwinska MW, James A, Devlin JT. Inferior Parietal Lobule Contributions to Visual Word Recognition. J Cogn Neurosci 2015; 27

Stevenson CM, Brookes MJ, Morris PG. β-Band correlates of the fMRI BOLD response. Hum Brain Mapp 2011; 32: 182–197.

Tarkiainen A, Helenius P, Hansen PC, Cornelissen PL, Salmelin R. Dynamics of letter string perception in the human occipitotemporal cortex. Brain 1999; 122: 2119–2132.

Taylor JSH, Rastle K, Davis MH. Can cognitive models explain brain activation during word and pseudoword reading? A meta-analysis of 36 neuroimaging studies. Psychol Bull 2013; 139: 766–791.

Thesen T, McDonald CR, Carlson C, Doyle W, Cash S, Sherfey J, et al. Sequential then interactive processing of letters and words in the left fusiform gyrus. Nat Commun 2012; 3: 1284–1288.

Twomey T, Kawabata Duncan KJ, Price CJ, Devlin JT. Top-down modulation of ventral occipito-temporal responses during visual word recognition. Neuroimage 2011; 55: 1242–1251.

Ueno T, Meteyard L, Hoffman P, Murayama K. The ventral anterior temporal lobe has a necessary role in exception word reading. Cereb Cortex 2018; 28: 3035–3045.

Vinckier F, Dehaene S, Jobert A, Dubus JP, Sigman M, Cohen L. Hierarchical Coding of Letter Strings in the Ventral Stream: Dissecting the Inner Organization of the Visual Word-Form System. Neuron 2007; 55: 143–156.

Wang L, Jensen O, Van den Brink D, Weder N, Schoffelen JM, Magyari L, et al. Beta oscillations relate to the N400m during language comprehension. Hum Brain Mapp 2012; 33: 2898–2912.

Watson CL, Possin K, Allen IE, Hubbard HI, Meyer M, Welch AE, et al. Visuospatial Functioning in the Primary Progressive Aphasias. J Int Neuropsychol Soc 2018; 24: 259–268.

Weekes BS. Differential effects of number of letters on word and nonword naming latency. Q J Exp Psychol Sect A Hum Exp Psychol 1997; 50: 439–456.

Wilkinson DT, Halligan PW, Marshall JC, Büchel C, Dolan RJ. Switching between the forest and the trees: Brain systems involved in local/global changed-level judgments. Neuroimage 2001; 13: 56–67.

Wilson M, Joubert S, Ferré P, Belleville S, Ansaldo A, Joanette Y, et al. The role of the left anterior temporal lobe in exception word reading: Reconciling patient and neuroimaging findings. Neuroimage 2012; 60: 2000–2007.

Wilson SM, Brambati SM, Henry RG, Handwerker DA, Agosta F, Miller BL, et al. The neural basis of surface dyslexia in semantic dementia. Brain 2009; 132: 71–86.

Woollams AM, Lambon-Ralph MA, Plaut DC, Patterson K. SD-squared: On the association between semantic dementia and surface dyslexia. Psychol Rev 2007; 114: 316–339.

Yuan H, Liu T, Szarkowski R, Rios C, Ashe J, He B. Negative covariation between task-related responses in alpha/beta-band activity and BOLD in human sensorimotor cortex: An EEG and fMRI study of motor imagery and movements. Neuroimage 2010; 49: 2596–2606.

